# A Framework for Comparing Mouse Neoantigen Immunogenicity

**DOI:** 10.64898/2026.02.10.704454

**Authors:** Peter J. Matulich, Carly N. Sprague, Victoria P. Schuster, Alyssa M. Granados, Rohan B. Chaudhari, Bree M. Mohr, Megan K. Ruhland, Megan L. Burger

## Abstract

Cytotoxic CD8+ T cell responses targeting tumor neoantigens are critical for immunotherapy efficacy and are widely studied across different preclinical mouse tumor models. Defined neoantigens are commonly identified or introduced into mouse tumors to enable tracking of tumor-specific T cell responses. However, these neoantigens differ in immunogenicity and variation in their composition may produce immune phenotypes that are mistakenly attributed to tumor-intrinsic mechanisms and complicate comparison across studies. Here, we sought to create a framework for comparing neoantigen immunogenicity by determining relative peptide-major histocompatibility complex (pMHC) affinity (K_D_) and stability (K_off_) and *in vivo* neoantigen-targeted vaccine responses for 25 commonly used mouse tumor-derived and model neoantigens. We found that *in silico* prediction of pMHC affinity moderately stratified strong versus weak *in vivo* immunogenicity but could not accurately predict rank order; experimental measurements of affinity did not improve prediction. In contrast, experimental measurement of pMHC stability closely correlated with the relative magnitude of vaccine responses demonstrating strong predictive value. Thus, we report the relative stability of commonly used neoantigens as a resource and provide a simple method to benchmark novel neoantigens against this library. This tool allows contextualization of the level of immunogenicity of newly identified neoantigens and aids in comparative interpretation of tumor-immune phenotypes across studies.

**SIGNIFICANCE:** Peptide-MHC stability strongly correlates with the in vivo response a neoantigen elicits, and this framework enables comparison of neoantigens across studies to evaluate the neoantigen contribution to the anti-tumor T cell response.

## INTRODUCTION

Immunotherapy has dramatically reshaped the treatment of many cancers, particularly in the metastatic setting. Immune checkpoint blockade (ICB) has had the most success in the clinic, leading to remarkable improvements in patient outcomes and fueling a surge of research into cancer immunology and immunotherapy^1^. As tumor cells accumulate mutations, a subset can generate an immune response. These immunogenic proteins, or neoantigens, are recognized as foreign by T cells and are the key mediators of ICB response^2–4^. A new generation of immunotherapies are being developed, including personalized cancer vaccines and adoptive T cell (ACT) therapies, that rely on identifying and targeting specific neoantigens^5^. Neoantigen-targeted ACT has achieved its first FDA approval in synovial sarcoma and personalized vaccines have reached late-stage clinical trials with a positive Phase III outcome in melanoma^6,7^. Understanding the regulation of neoantigen-specific T cell responses, including which mutations can best be leveraged therapeutically, will be critical for increasing the impact of these approaches.

While studying neoantigen-specific T cells is a clear priority, identifying them in patient samples presents significant challenges. Neoantigens are unique to each patient, requiring exome sequencing of individual tumor samples; computational algorithms then predict tens to hundreds of possibly immunogenic epitopes. To validate predicted neoantigens, tumor-infiltrating T cells must be directly tested for neoantigen-reactivity^2–4^ and often only a single digit number elicit a detectable response. Given this labor-intensive process, investigators have primarily used mouse models expressing defined neoantigens to study neoantigen-specific T cell responses, which have been invaluable tools for understanding tumor-immune interactions^8,9^. Typically, tumor-specific CD8+ or CD4+ T cell responses are studied by either expressing a model neoantigen (e.g., SIINFEKL from chicken ovalbumin) in a tumor or using a tumor cell line with known neoantigens (e.g., MC38 colon adenocarcinoma). The T cell responses to these neoantigens are then evaluated by monitoring either the endogenous T cell response or transferred transgenic T cells with a neoantigen-specific T cell receptor (TCR). It is increasingly clear that the context and identity of neoantigen expression can significantly influence the anti-tumor T cell response. Antigen immunodominance hierarchies form wherein the T cell response is dominated by the most immunogenic neoantigen, while responses against weaker neoantigens are suppressed^10,11^

Therefore, ranking neoantigens is critical for accurately quantifying the magnitude of the tumor-specific T cell response, as contribution can dramatically shift between the strongest and second-strongest antigen in a system. Some antigens are also intrinsically more or less immunogenic and can considerably reshape outcome in terms of tumor control. For example, the neoantigen Mrps35 was recently shown to generate a CD8+ T cell response in the common KPC 4662 pancreatic cancer cell line^12^; while these cells readily form tumors, engineering in Ova containing the model neoantigen SIINFEKL leads to spontaneous rejection^13^. Developing a way to compare neoantigen immunogenicity across studies is therefore essential to disentangle neoantigen-intrinsic from tumor-intrinsic phenotypes.

Several factors are known to influence neoantigen immunogenicity, with the primary determinant being the density of peptide-MHC (pMHC) complexes at the cell surface^14–16^. Because empty MHC molecules are unstable, a neoantigen’s ability to load onto MHC and remain bound is critical for determining its surface density. Antigen loading is tied to binding affinity, while the persistence of the pMHC complex at the cell surface is governed by its stability^15,17^. *In silico* algorithms have been developed to predict immunogenic peptides and their binding affinity^18,19^. These algorithms are trained on large datasets of experimentally measured pMHC affinities and large-scale peptide elution studies in which MHC-bound peptides are identified by mass spectrometry^18,19^. As these algorithms are trained in part on binding affinity, they can report a predicted binding affinity for antigens. To date, benchmarking efforts have largely evaluated these algorithms on their ability to separate binders from non-binders, using classification metrics such as the area under the receiver operating characteristic curve (AUC)^20–22^. However, it remains unclear how effectively current prediction algorithms can discern relative immunogenicity among a group of mouse neoantigens and whether these algorithms can confidently be used to compare neoantigens across studies, representing a key gap that we aim to address.

In this study, we selected 19 commonly studied mouse tumor-derived neoantigens^23–31^ and 6 model neoantigens^32–35^ from various tumor systems and evaluated the concordance of *in silico* prediction outcomes with experimental measures of relative pMHC affinity and stability. We further assessed alignment of both *in silico* and experimental measures with ground-truth immunogenicity in an *in vivo* vaccine setting. We found that *in silico* predictions moderately stratified strong versus weak *in vivo* immunogenicity of the neoantigens, but were unable to accurately predict rank order, especially for strong neoantigens. Interestingly, experimental pMHC binding affinity measurements exhibited a worse correlation; however, experimental pMHC stability measurements were highly correlated with *in vivo* immunogenicity, consistent with accumulating evidence supporting stronger predictive value of stability versus affinity in recent studies^11,17,36–38^. These findings nominate the experimental measurement of pMHC stability as a strong metric to compare neoantigens across studies. This work defines the relative pMHC stability of a set of commonly used mouse neoantigens as a resource for the field and provides a straightforward framework and tool for benchmarking novel neoantigens against this reference to aid in comparison of neoantigen-driven immunity across tumor models.

## RESULTS

### *In Silico* Algorithms Similarly Rank the MHC I Binding Affinity of Mouse Neoantigens

We sought to computationally predict and experimentally determine the relative immunogenicity of a library of 25 CD8+ T cell-reactive neoantigens. We selected 19 mouse tumor-derived neoantigens from seven common tumor systems, including MC38 colon adenocarcinoma and methylcholanthrene (MCA)-induced sarcoma, and added six commonly used model neoantigens, including antigens from chicken Ovalbumin and lymphocytic choriomeningitis virus (LCMV) (**Fig. 1A**). We first passed the assembled neoantigen peptide sequences through nine publicly available *in silico* prediction algorithms that support the mouse MHC-I alleles: NetMHCpan4.2^39^, NetMHCpan4.1^40^, NetMHCpan4.0^18^, ANN4.0^41^, SMMPMBEC 1.0^42^, SMM1.0^43^, PickPocket1.1^44^, NetMHCcons1.1^45^ and MHCflurry2.0^19^. Most of these algorithms are trained as pan-allele models, allowing them to generate predictions for mouse alleles even when human data were the primary training focus, with ANN, SMM and SMMPMBEC being exceptions using only data available for the specific allele. Most methods rely on neural networks trained on pMHC binding affinity data and MHC elution mass spectrometry datasets. NetMHCcons1.1 is a consensus model that integrates predictions from NetMHCpan2.8, ANN3.4 and PickPocket1.1 to improve overall performance. These algorithms predict the binding affinity, or K_D_, of peptides for their cognate MHC-I molecule (**Fig. 1A**) reported as an IC50, or the concentration of peptide required to bind 50% of the available MHC molecules. Our neoantigens were assigned a wide range of predicted affinities by these algorithms, ranging from 1.83 nM to 6469 nM. Generally, neoantigens with a binding affinity of >500 nM are considered unlikely to elicit a CD8+ T cell response, while a binding affinity of <50 nM predicts strong immunogenicity^14^. When applying these criteria, the majority of programs classified most of the neoantigens as immunogenic, with the notable exception of PickPocket1.1, which classified only 60% of the neoantigens below 500 nM and 13% below 50 nM. At the other extreme, MHCflurry classified 80% of the neoantigens as strong (<50 nM), substantially more than the next highest predictor, NetMHCpan4.1, at 60%. Given every neoantigen in our library was originally identified by its ability to elicit a CD8+ T cell response *in vivo*, these findings suggest PickPocket1.1 is a poor predictor of immunogenicity for mouse neoantigens, and the broad predicted affinity values across algorithms highlights the limitations of these models.

**Figure 1.**
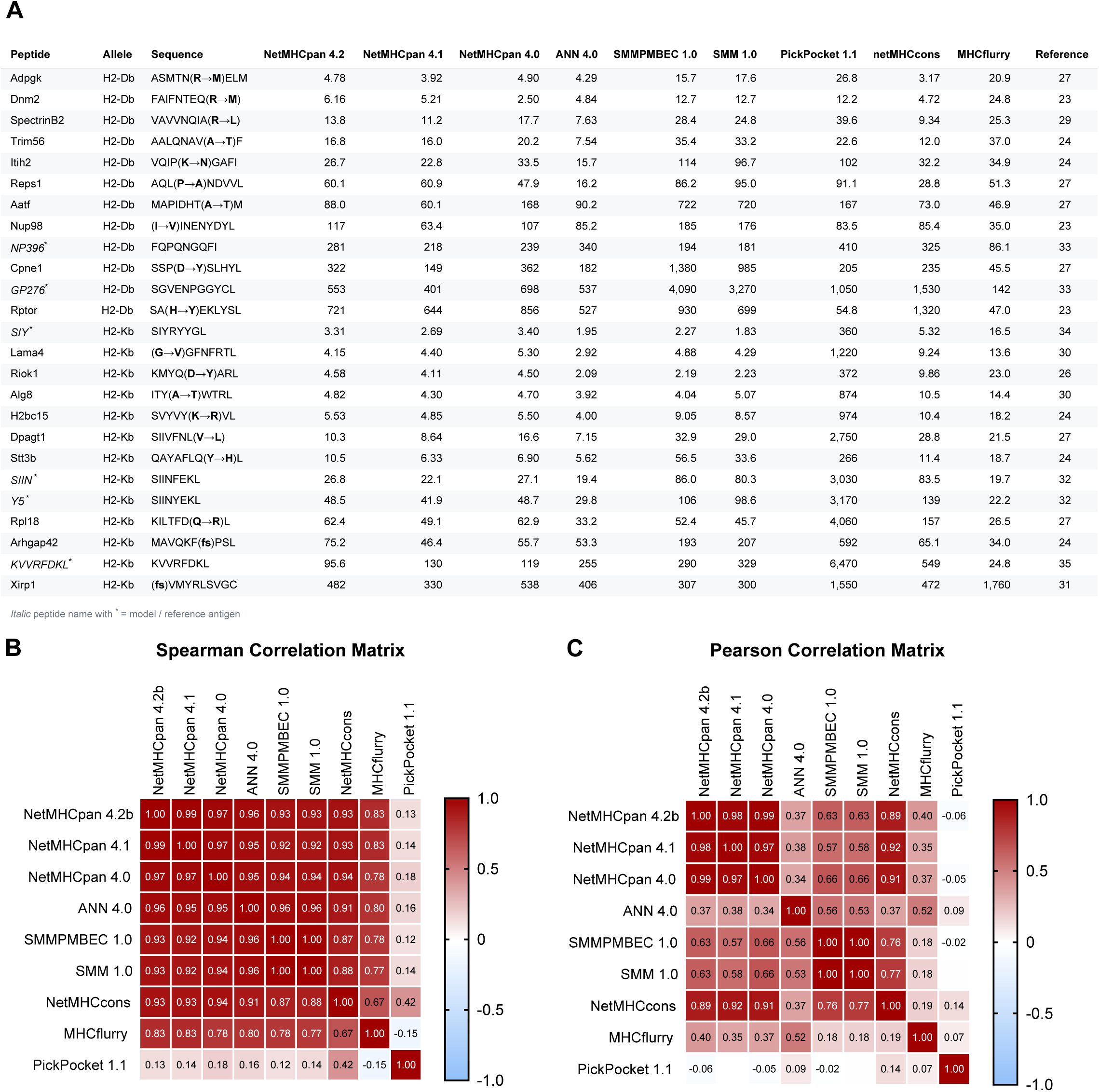
In Silico Algorithms Similarly Rank the MHC I Binding Affinity of Mouse Neoantigens. (A) Table of mouse antigens with tumor-derived neoantigens annotated with single nucleotide variant or frameshift denoted as (fs) and model antigens marked with italics and *. C57BL/6 MHC allelic restriction (H2-D^b^ or H2-K^b^), reference and predicted IC50 values from the *in silico* algorithms are indicated. (B-C) Correlation matrix of the predicted IC50 values for the neoantigen library in (A) between different *in silico* algorithms using the Spearman correlation coefficient (B) or Pearson correlation coefficient (C).

The algorithms generally ranked the neoantigens in a similar order with high Spearman correlation coefficients across both alleles tested (**Fig. 1B**). PickPocket 1.1 was an outlier, showing little correlation with the other models, likely driven by its consistently weaker MHC affinity predictions. MHCflurry was also less correlated with the other algorithms and generally predicted higher affinities. Whereas the Spearman coefficient captures how consistently the algorithms rank neoantigens in the same order, the Pearson coefficient reflects the strength of the linear relationship between their predicted affinity values. When using the Pearson correlation coefficient to compare the linear relationship of binding affinities across the algorithms, the outputs generally grouped based on commonalities in the training data for the algorithms, with the pan-allele algorithms grouping separately from the allele-specific algorithms (**Fig. 1C**). Overall, the Spearman correlation coefficients were more closely aligned across the models than the Pearson, indicating that there is stronger agreement in the rank order of antigens than absolute affinity values.

### *In Vitro* Affinity Measurements Poorly Correlate with *In Silico* Values

To determine how predicted binding affinities align with experimentally determined values, we employed an RMA-S peptide affinity assay^46^, which is commonly used to experimentally determine the pMHC equilibrium dissociation constant (K_D_). RMA-S cells, which produce empty MHC I molecules, are incubated with a range of exogenous peptide concentrations that stabilize the MHC molecules at the cell surface. The stabilized MHC is then quantified by antibody staining and flow cytometry, with the relative K_D_ defined as the peptide concentration occupying 50% of surface MHC. As the assay utilizes allele specific antibodies, it provides a relative measure of pMHC binding that is best compared within a given allele. Performing this assay on our library of neoantigens, we observed a range of K_D_ values spanning several orders of magnitude (**Fig. 2A-2B**; neoantigens listed in order of ascending K_D_ value). For the neoantigens binding the H2-K^b^ allele, the strongest binder was SIINFEKL, which is the most used model neoantigen across mouse tumor studies, followed closely by the tumor-derived neoantigen H2bc15 from the Kras/p53-driven genetically engineered mouse model of lung adenocarcinoma^24^. The other model neoantigens exhibited K_D_ values within the range of tumor-derived neoantigens. These included SIYRYYGL (SIY), a common model antigen recognized by 2C T cells ^34^, and SIINYEKL (Y5), a weaker MHC-binding variant of SIINFEKL that is recognized by OT-I T cells^32^. Neoantigens binding the H2-D^b^ allele displayed a more compact range of K_D_ values, with immunodominant LCMV-derived NP396 narrowly assuming the top position. Only two neoantigens exhibited a log-fold weaker K_D_ than the top antigen, and both were predicted by the majority of the *in silico* algorithms to be among the weaker binders in the library. However, notably, the strongest binder, NP396, was also predicted to be among the weakest binders by most of the *in silico* algorithms.

**Figure 2.**
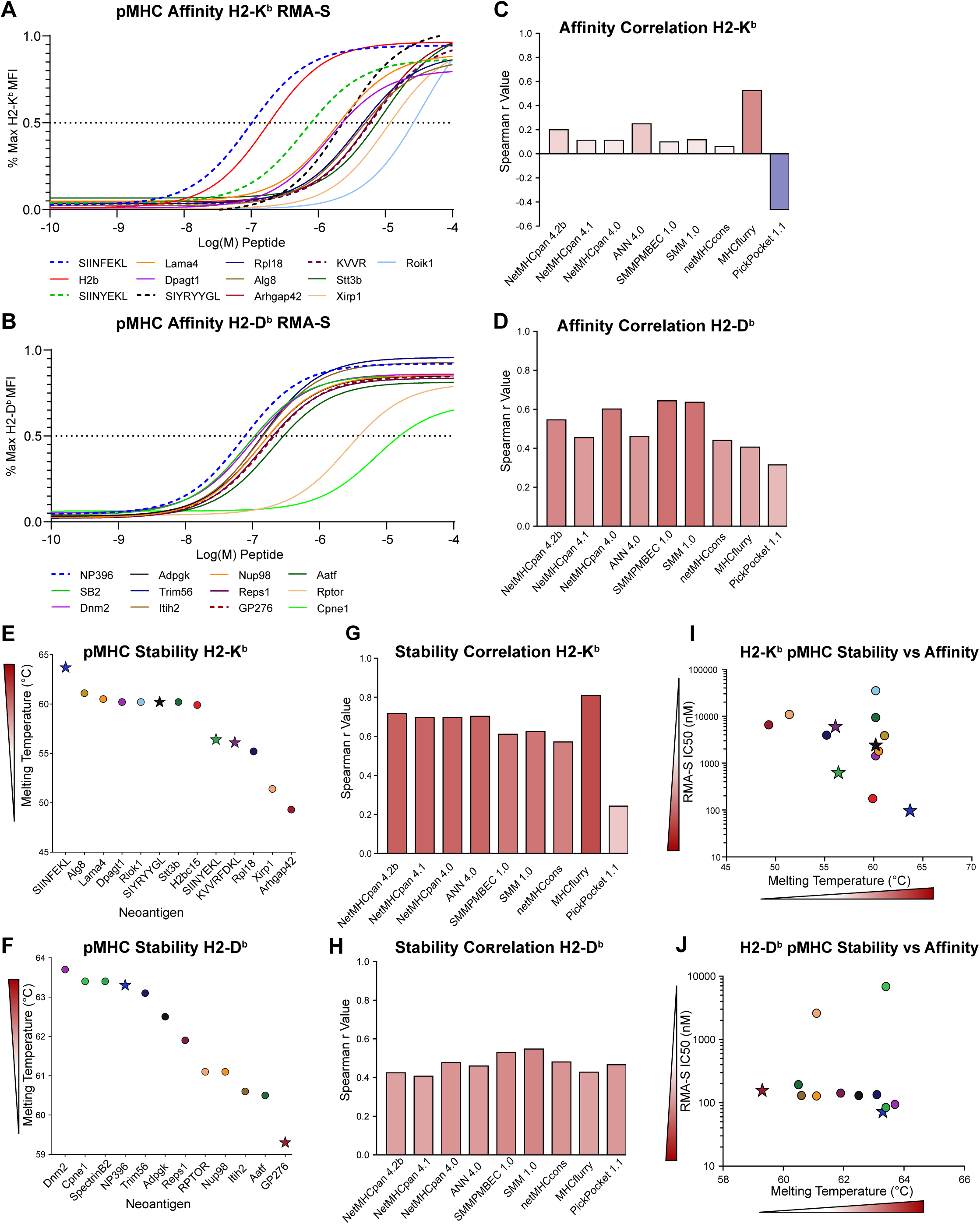
In Silico Predictions Reflect In Vitro Stability More Than Affinity. (A-B) Relative peptide-MHC (pMHC) affinity (K_D_) for each neoantigen as assessed by flow cytometry staining for stabilized MHC-I molecules, H2-K^b^ (A) and H2-D^b^ (B), on RMA-S cells incubated with exogenous peptide. The plotted curve uses an S4PL model with a Hill Slope of 1. Legends place the neoantigens in order of ascending K_D_. Data are from a representative experiment of n = 2-3 independent experiments, each with 3 technical replicates. (C-D) Plotted Spearman correlation coefficients for experimentally determined (RMA-S assay) versus *in silico* predicted IC50 values for H2-K^b^ (C) and H2-D^b^ (D). (E-F) Relative pMHC stability (K_off_) for each neoantigen as measured by differential scanning fluorimetry of H2-Kb (E) and H2-D^b^ (F) pMHC thermal melting. Tumor-derived neoantigens are denoted as circles; model neoantigens as stars. Melting temperature of the complex is defined as the minimum value of the inverse derivative of fluorescence over time (-dF/dT). Wedge indicates relative strength of neoantigen. Data are from a representative experiment of n = 2-3 independent experiments, each with 3 technical replicates. (G-H) Plotted Spearman correlation coefficients for experimentally determined stability versus *in silico* predicted IC50 values for H2-K^b^ (G) and H2-D^b^ (H). (I-J) RMA-S assay pMHC affinity IC50 values plotted against the pMHC stability melting temperature values for each neoantigen for the H2-K^b^ (I) and H2-D^b^ (J) alleles. Wedge indicates relative strength of neoantigen.

In general, there was poor agreement between the ranking of neoantigen affinities by the *in silico* models and the experimentally measured K_D_ values, with low Spearman correlation coefficients and no models passing a Spearman correlation of 0.6 (**Fig. 2C-2D**). The models performed better at ranking neoantigens that bind the H2-D^b^ allele than the H2-K^b^ allele; however, the strongest correlation coefficient across either allele was 0.54. The disagreement seemed most pronounced among the strongest binding neoantigens, which the algorithms consistently ranked poorly relative to their measured affinities. Collectively, these findings suggest that *in silico* prediction programs can generally predict MHC binders and approximate binding affinity but poorly rank the MHC binding affinities among neoantigens.

### pMHC Stability Poorly Correlates with Affinity

Another measurement of MHC binding strength is the dissociation rate of the pMHC complex, K_off_ or stability, which is related to affinity (K_D_) as K_D_ = K_off_/K_on_. The melting temperature of pMHC is inversely proportional to K_off_, where a higher melting temperature is equivalent to a longer half-life of the pMHC complex^47^. Multiple studies suggest stability can better predict the immunogenicity of an antigen than affinity in both viral and tumor models^11,17,36–38^, despite the wider use of affinity as a prediction metric. We used UV-exchangeable MHC monomers to generate pMHC complexes for our neoantigen library and generated pMHC melting curves by differential scanning fluorimetry^47^. In line with our pMHC affinity measurements, SIINFEKL had a higher melting temperature (indicative of a lower K_off_) than the other H2-K^b^ neoantigens (**Fig. 2E-2F**). Again, neoantigens binding the H2-K^b^ allele displayed a wider range of values compared to H2-D^b^, with greater representation of weaker binders. Melting temperature values generally correlated better with the *in silico* predicted affinities than the experimentally measured pMHC binding affinities did, which was somewhat surprising given the *in silico* algorithms predict K_D_ and not K_off_ (**Fig. 2G-2H**). MHCflurry2.0 showed the highest correlation with both the experimentally measured pMHC affinity and stability for the H2-K^b^ allele with a Spearman correlation coefficient of stability reaching 0.81 representing a strong correlation. PickPocket1.1 showed a markedly weaker correlation than the other algorithms across both affinity and stability, particularly for the H2-K^b^ allele, where there was a negative correlation. In contrast, for the H2-D^b^ allele, all *in silico* algorithms performed similarly for predicting stability, but no model exceeded a correlation coefficient of 0.6.

When comparing the rank order of neoantigens measured by the affinity and stability assays, there was some agreement for neoantigens binding the H2-K^b^ allele but little for H2-D^b^ (**Fig. 2I-2J**). A narrower range of melting temperatures for H2-D^b^ binders may have contributed, spanning only 4.4°C compared to 14.4°C for H2-K^b^ binders, and melting temperatures notably did not identify the two outlier H2-D^b^ neoantigens in the RMA-S assay. In contrast, neoantigens binding H2-K^b^ showed more of a trend between the stability and affinity measurements, and in cases where disagreement was observed, the neoantigens exhibited higher melting temperatures than would be predicted by their K_D_ values. Notably, these two measurements were not aligned suggesting these biochemical constants cannot be used interchangeably (**Fig. 2I-2J**).

### pMHC Stability Correlates with In Vivo Immunogenicity

To determine which measurements of pMHC binding strength best correlate with *in vivo* neoantigen immunogenicity, we vaccinated wild-type C57BL/6 mice individually with each of the neoantigens from our library. Mice were vaccinated with bone marrow-derived dendritic cells pulsed with a single neoantigen, followed by a single boost one week later consisting of a corresponding short-peptide vaccine formulated with a cyclic di-GMP adjuvant. The goal of our vaccine strategy was to elicit a below saturating response to achieve a dynamic range of responses for the neoantigen set; consistently, we observed detectable responses for the majority but not all of the neoantigens (**Fig. 3A-3B**). Overall, the *in silico* programs moderately predicted the rank order of the CD8+ T cell response to each neoantigen and largely no one algorithm outperformed the others (**Fig. 3C-3D**). Surprisingly, experimentally measured pMHC affinity was a worse predictor of *in vivo* immunogenicity (**Fig. 3E-3F**). In contrast, pMHC stability was the strongest predictor and accurately predicted rank order of the neoantigens, with Spearman correlation coefficients of 0.81 for the H2-K^b^ allele and 0.94 for H2-D^b^ (**Fig. 3G-3H**). These results demonstrate that while *in silico* prediction algorithms can moderately stratify strong versus weak neoantigens, *in vitro* measurement of pMHC stability has greater power to rank relative neoantigen immunogenicity *in vivo*.

**Figure 3.**
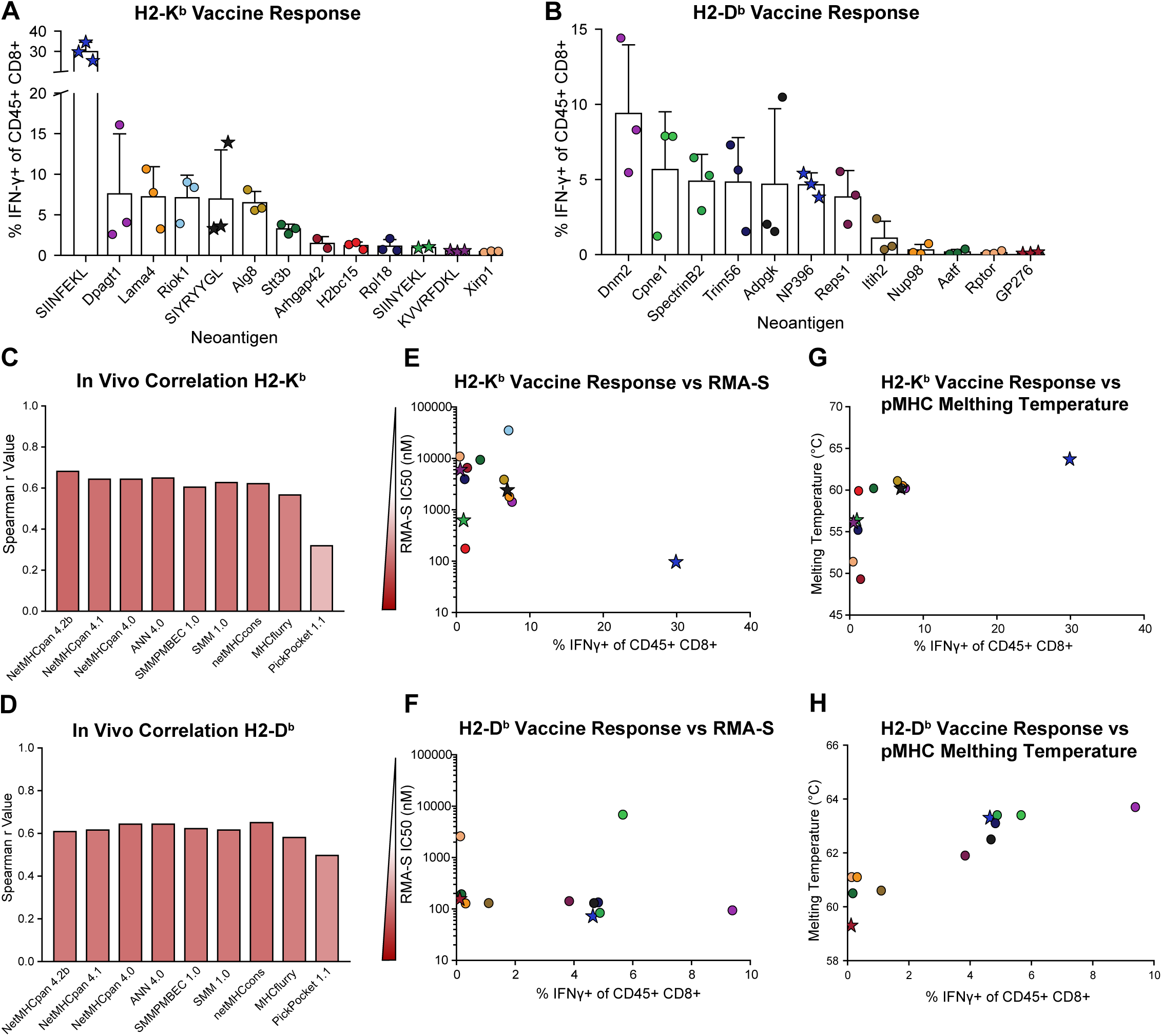
pMHC Stability Correlates with In Vivo Immunogenicity. (A-B) The percent of IFNγ+ CD45+ CD8+ T cells is graphed for each neoantigen-targeted vaccine response. Neoantigen peptide-pulsed bone marrow-derived dendritic cell vaccines (1.5 million cells) were delivered into wild-type C57BL/6 mice followed one week later by a booster consisting of neoantigen peptide (6 µg) with cyclic di-GMP adjuvant (25 µg). Data are from a representative experiment of n = 1-2 independent experiments, with 3 mice per group. (C-D) The plotted Spearman correlation coefficient for the percent IFNγ+ cells for each neoantigen in A and B compared to the predicted IC50 values from each *in silico* algorithm for the H2-K^b^ (C) and H2-D^b^ (D) alleles. (E-F) RMA-S assay pMHC affinity IC50 values plotted against the percent IFNγ+ cells for each neoantigen for the H2-K^b^ (E) and H2-D^b^ (F) alleles. Wedge indicates relative strength of neoantigen. (G-H) Plot of pMHC stability melting temperature values against the percent IFNγ+ cells for each neoantigen for the H2-K^b^ (G) and H2-D^b^ (H) alleles.

### Training Datasets Poorly Rank Strong Neoantigens

A potential source of the error in *in* silico ranking of *in vivo* immunogenicity could be the training data, where discordance between measured affinities has been observed among strong antigens. A benchmarking study comparing affinity measurements made in two independent laboratories by different methods found that measurements were mostly reproducible, except among the strongest antigens^48^. We investigated whether the same observation could be made in the large training datasets used by the majority of prediction algorithms. MHCflurry and NetMHCpan, the two most commonly used algorithms, both draw on data from the Immune Epitope Database (IEDB), a public database that acts as a catalog of experimental data of T and B cell epitopes^49^. Using the IEDB dataset used to train the MHCflurry model^19^, we examined antigens with more than one experimental measurement and assessed the level of agreement between measurements. Of the 713,069 entries, there were 4,174 pairs of repeated quantitative affinity measurements. Pairs were plotted against each other and measurements correlated (**Fig. 4A-4B**).

**Figure 4.**
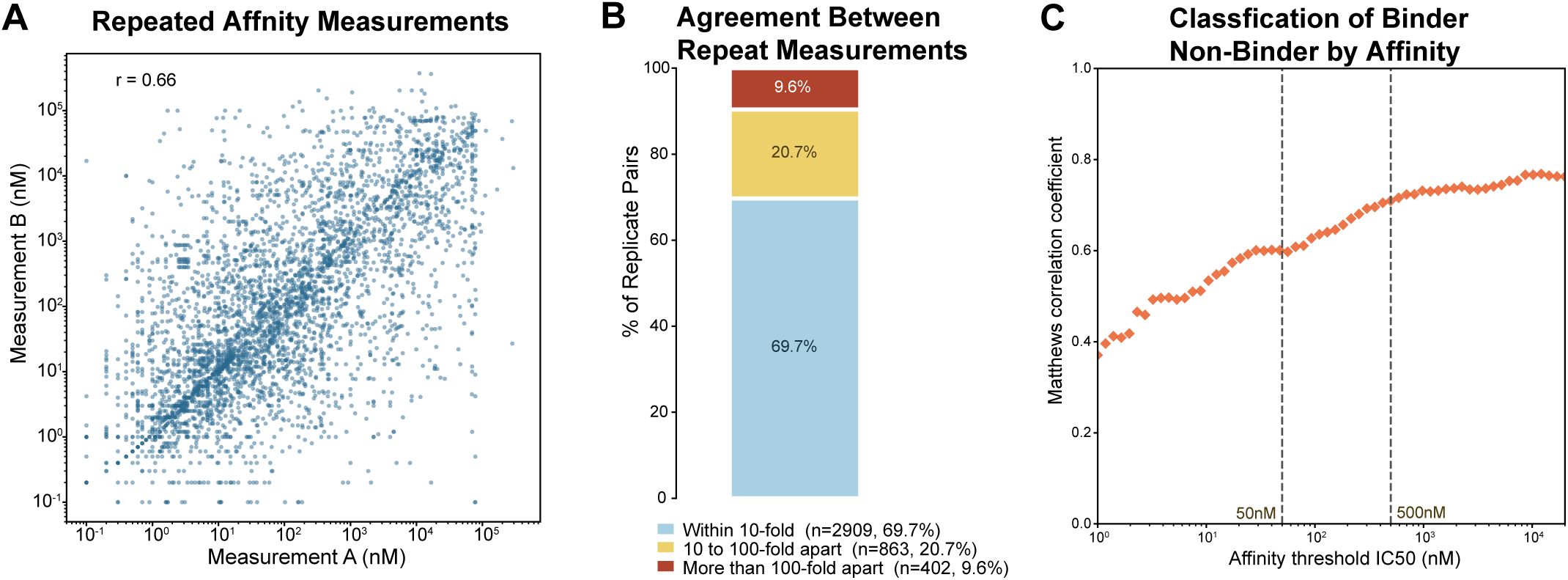
Training Datasets Poorly Rank Strong Neoantigens. (A) Scatter plot comparing replicate measured affinities for IEDB antigens included in the MHCflurry training dataset. (B) Distribution of the replicate pairs in A (4,174 total) by the magnitude of their disagreement. (C) Matthews correlation coefficient (MCC) as a function of the IC₅₀ threshold used to classify peptides as binders or nonbinders. Each point represents the MCC of the classification of binder/non-binder for the replicate pairs at that threshold. The dashed lines mark IC₅₀ = 50 nM and 500 nM as standard immunogenicity cutoffs.

At scale, repeat measurements correlated reasonably well (Pearson correlation coefficient of 0.66); however, 30.3% of the pairs differed by more than an order of magnitude, and 9.6% differed by more than two orders of magnitude (**Fig. 4B**). To assess agreement across the range of affinities, we computed the Matthews correlation coefficient between entries over a sliding scale of threshold values (**Fig. 4C**). Agreement was poorest among the strongest antigens: the Matthews correlation coefficient was lowest below 50 nM and climbed from ∼0.4 to ∼0.7 as affinities weakened toward the 500 nM immunogenicity threshold. This zone of greatest disagreement coincides with the strongest antigens that the prediction algorithms ranked poorly throughout our dataset. This underlying irreproducibility and noise among the strongest antigens in the training data may limit the ability of *in silico* prediction algorithms to accurately rank strong antigens.

### pMHC Stability Measurement to Benchmark Immunogenicity of Novel Neoantigens

Our observations that pMHC stability strongly predicts *in vivo* immunogenicity align with publications demonstrating stability to be a more accurate measurement of immunogenicity than affinity across viral and tumor contexts^11,17,36–38^. The melting temperature assay we used to measure stability can be easily replicated and may aid investigators in better estimating the relative immunogenicity of novel neoantigens in their systems. MHC-I monomers are currently available to academic investigators through the NIH Tetramer Core free of charge and the melting temperature assay can be performed with a standard real-time PCR machine, making measurement of pMHC stability a simple and low-cost method for assessing neoantigen immunogenicity. We propose benchmarking novel neoantigens against a known strong and weak neoantigen from our library as standards to facilitate comparison of neoantigen immunogenicity across labs and reagent lots; specifically, SIINFEKL and Y5 as a pair for H2-K^b^ and NP396 and GP276 as the pair for H2-D^b^. By measuring the melting temperature of these two standards, one can determine the strength of an unknown. To express relative strength on an arbitrary scale anchored to two references, SIINFEKL is set to 100 and Y5 to 0. Relative strength for a novel peptide is then calculated as:

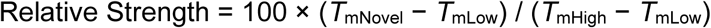

where *T*_mNovel_, *T*_mHigh_, and *T*_mLow_ are the melting temperatures of the test peptide and the strong and weak standards, respectively.

As an example, we evaluated additional H2-K^b^-restricted neoantigens not originally in the library, GP34^33^, Trp2^50^, Med12^51^, Yipf4^25^, and Mrps35^12^, and plotted their melting temperature against SIINFEKL and Y5 standards to determine their relative strength (**Fig. 5A**). The strength of the neoantigens was then graphed against the H-2K^b^ subset of the neoantigen library (**Fig. 5B**). We observed that the new neoantigens were on the low end of the range, with GP34, a subdominant LCMV neoantigen, nearest the H2bc15 epitope in strength, while the tumor-associated melanoma neoantigen Trp2^50^ and the Mrps35 neoantigen from KPC 4662 pancreatic cancer^12^ were approximately the same strength as Y5. To assist users in performing similar comparisons, we have created a freely available web tool (available at https://pmhc-stability-benchmark.web.app/) that takes a user-submitted pMHC melting temperature for a novel neoantigen and two standards as input and returns the relative strength of the novel neoantigen. The tool is maintained by the Burger Lab and will be updated as additional neoantigens are benchmarked by our lab. Additionally, through the web tool, users can submit their own measurements to be added to the library as a community resource. As this reference set grows and new neoantigens across different tumor models are entered, we anticipate it will increase in value, allowing for improved contextualization of immunogenicity across studies and models.

**Figure 5.**
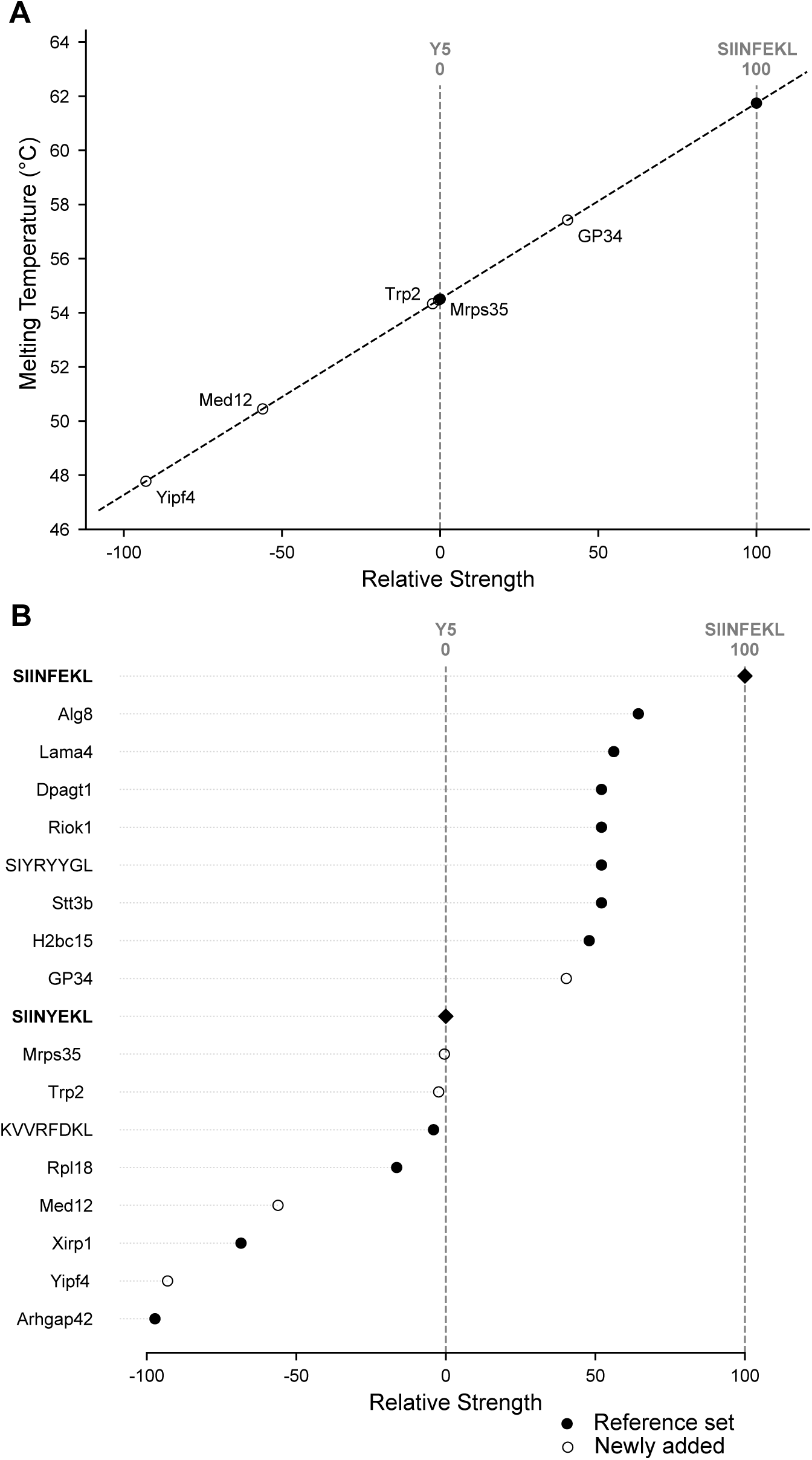
pMHC Stability Measurement to Benchmark Immunogenicity of Novel Neoantigens. (A) Plot of melting temperatures measured by differential scanning fluorimetry for the H-2K^b^ model and tumor neoantigens GP34, Trp2, Med12, Yipf4, and Mrps35 using SIINFEKL and SIINYEKL as benchmarks to determine their relative strength. (B) Plot of the relative strength of the neoantigens from A (empty circles) against the other H2-Kb neoantigens in the library in Fig. 1A (closed circles).

## DISCUSSION

A persistent challenge in the preclinical study of tumor neoantigens is the lack of a reliable way to compare neoantigen strength across tumor models; this makes it difficult to compare results across studies, as anti-tumor T cell responses are shaped by both tumor-intrinsic factors and neoantigen-intrinsic properties. In this work, we have shown that a simple *in vitro* approach of measuring pMHC stability can benchmark relative neoantigen immunogenicity and provide an important framework for comparing neoantigen-driven phenotypes across studies.

The importance of such a metric is underscored by how strongly neoantigen identity can shape anti-tumor immunity. As discussed above, the KPC 4662 pancreatic cancer model harbors an immunogenic endogenous neoantigen, but this neoantigen does not prevent tumor outgrowth; however, engineering in the strong model neoantigen SIINFEKL drives spontaneous tumor rejection, demonstrating that the strength, and not merely the presence, of a neoantigen governs the response^13^. Furthermore, a neoantigen may drive tumor rejection in one context, while allowing tumor outgrowth in another; for example, Spectrin-β2 drives rejection of MCA-induced sarcoma cell line d42m1, but not KP9025 Kras/p53-driven sarcoma^52^. The complexity of the comparison further increases when both the tumor context and neoantigen repertoire are not conserved, as is routinely the case in therapeutic studies. For example, an M13 bacteriophage vaccine against SIINFEKL in B16-Ova was more efficacious than against Adpgk in MC38; better tumor control of B16-Ova may have been attributable to the relatively greater immunogenicity of SIINFEKL rather than tumor-intrinsic differences, particularly given MC38 is typically more immunotherapy-responsive^53^. Hence, placing commonly studied mouse neoantigens on a relative-strength scale will allow better attribution of differential phenotypes across studies to neoantigen-intrinsic versus tumor-intrinsic differences.

This study evaluated the extent to which *in silico* predictions and *in vitro* measurements of pMHC binding strength could stratify *in vivo* immunogenicity of a library of known mouse tumor and model neoantigens. We found that *in silico* algorithms performed worse than *in vitro* pMHC stability measurements at ranking the neoantigens in order of *in vivo* immunogenicity. The *in* silico algorithms may perform suboptimally because these algorithms were not trained to discern the difference in strength between immunogenic antigens, but rather to select the peptides that are likely to bind MHC out of a much larger set of non-binders. In line with this, our meta-analysis of peptides used to train MHCflurry with multiple MHC affinity measurements showed that measurements for the subset of antigens with the highest binding affinities (<50 nM) were poorly correlated and the peptide rank order was irreproducible. These algorithms are also trained in large part on eluted peptide quantification data, which may overrepresent more stable peptides; this could explain why their predictions correlated better with our measured pMHC stability than with measured affinity, despite their output being predicted affinity. Ultimately, this suggests *in silico* prediction algorithms are inadequately equipped to accurately rank strong neoantigens due to the noise in the underpinning training data.

A growing body of research suggests that pMHC stability is more predictive of antigen immunogenicity than pMHC affinity. In a study comparing immune responses to vaccina virus antigens, pairs of antigens with the same affinity were found to generate sizably different CD8+ T cell responses; the difference was attributed to differential stability, where the more stable antigen consistently garnered the largest immune response^17^. Only recently has this been extended to tumor neoantigens, where we and others found that the relative stability, rather than affinity, of neoantigens can predict the identity of neoantigens capable of driving neoantigen-specific CD8+ T cell responses^11,51,54^. This also appears to hold true in the context of therapeutic vaccination, where an mRNA-lipid nanoparticle vaccine concurrently expressing multiple neoantigens elicited the largest CD8+ T cell responses against the highest stability neoantigens^51^. Consistent with these observations, experimental measurement of relative pMHC stability by differential scanning fluorimetry accurately predicted the relative *in vivo* immunogenicity of our library of neoantigens. Therefore, we propose that pMHC stability is a reliable metric to predict the relative strength of neoantigens and have provided a framework for benchmarking newly identified mouse neoantigens against our reference set using a simple pMHC melting temperature-based assay, implemented as a freely available web tool (https://pmhc-stability-benchmark.web.app/).

In the absence of well-characterized tumor-derived neoantigens, many studies have relied on model neoantigens, such as SIINFEKL and others included here, as surrogates to evaluate tumor neoantigen-specific T cell responses. Model neoantigens offer several advantages, including generally strong immunogenicity, decades of use across multiple tumor models and the wide availability of experimental tools for tracking neoantigen-specific T cell responses, including pMHC tetramers and T cell receptor (TCR) transgenic mice. However, it has been theorized that immunoediting may bias tumors toward a repertoire of weaker antigens, potentially limiting the relevance of model neoantigens for preclinical study^55^. In our study, we did not observe a broad distinction between tumor-derived neoantigens and commonly used model neoantigens in terms of pMHC binding strength or *in vivo* vaccine response. This is consistent with several studies showing that human tumors can harbor mutations with very strong neoantigens, including with higher affinity than SIINFEKL^56^. Notably, two recent studies in genetically engineered mouse models of lung and pancreatic cancer also observed that CD8+ T cell responses against SIINFEKL and the MCA-induced sarcoma neoantigens Alg8 and Lama4 were of comparable magnitude and phenotype^11,57^. In our dataset, Alg8 and Lama4 ranked near but lower than SIINFEKL in relative pMHC stability; these findings might suggest that in a given tumor setting, there may be a threshold of stability that yields a maximal T cell response.

Despite the strong predictive value of pMHC binding stability, this framework has limitations. While pMHC stability is a major determinant of surface pMHC density, baseline expression of the protein also influences antigen presentation. For example, in MC38 colon adenocarcinoma, Rpl18 has been reported to be immunodominant^28^, despite relatively low pMHC affinity and stability; however, this antigen has high expression as a ribosomal protein, with its transcript roughly 30-fold more abundant than that of the next most dominant neoantigen, Adpgk^58^. This underscores that relative pMHC stability does not capture antigen expression level or the presence of low-frequency subclonal variants, both of which can also shape immune responses. High pMHC stability may also not predict greater immunogenicity if pMHC affinity is dramatically lower. In a Kras/p53-driven model of lung adenocarcinoma where six individually-expressed neoantigens were compared (Adpgk, SpectrinB2, Lama4, Alg8, Aatf, and Cpne1)^10^, our relative stability measurements would have correctly predicted the *in vivo* responses of five. The sixth, CpneI, had high stability, but much lower affinity—orders of magnitude below the others—which may lead to poor peptide loading *in vivo* that reduces its immunogenicity. Hence, pMHC stability is not a fail-proof predictor of immunogenicity and integration of other measurements including affinity or expression could improve accuracy. Additional factors have also been recognized to contribute to immunogenicity, including proteasomal processing^59–61^, naïve T cell precursor repertoire and TCR affinity for pMHC^62^.

As neoantigen prediction moves into the clinic with personalized cancer vaccines, now in Phase III trials^6^, the quality of neoantigen selection is increasingly important^63^. Studies of endogenous neoantigen responses consistently show that the number of predicted neoantigens far exceeds the number that elicit detectable T cell responses, and that the magnitude of these responses varies dramatically, ranging from only a few cells to thousands^2–4,64,65^. This narrowing of immune responses from predicted to observed is also true in neoantigen vaccine trials where often tens of neoantigens are vaccinated against but very few generate a detectable response^63,66–69^. Breadth may be particularly important for controlling clonally heterogeneous tumors, where responses directed at multiple antigens can prevent immune escape via immunoediting^55^. Our observations here suggest that future vaccine design may benefit from incorporating measured or predicted stability of pMHC complexes to assist in including the strongest epitopes and generating the maximum number of possible responses. While directly measuring pMHC stability may be challenging to incorporate into the clinical vaccine pipeline, it should be feasible to generate a large reference dataset of neoantigen stability measurements to train the next generation of prediction models for neoantigen selection. Ultimately, as neoantigen-targeted immunotherapies become an increasingly common tool in the fight against cancer, a deeper understanding of which neoantigens elicit immune responses and which can be leveraged to generate *de novo* responses is critical to maximizing their utility.

## METHODS

### RMA-S Cell MHC Affinity Assay

RMA-S cells were cultured in RPMI 1640 media (10% fetal bovine serum (FBS), 2 mM L-glutamine, 10 mM HEPES, 1X NEAA, 1X β-mercaptoethanol and 50 U/µL Penicillin/Streptomycin) at 37°C at 5% CO_2_ and transferred to 26°C at 5% CO_2_ for 4 hours to stabilize MHC molecules on the cell surface. Cells were then plated at 100,000 cells per well in 96-well flat bottom plates with 10-fold serial dilutions of the respective peptide for 1 hour at 26°C. Cells were moved to 37°C for one hour to internalize empty MHC molecules and then kept on ice to stop MHC trafficking. Cells were washed with FACS buffer (1% heat-inactivated FBS in PBS) and stained with either H2-K^b^ or H2-D^b^ antibody (Cat. No.17-5958-82, RRID: AB_1311280, Cat. No. 12-5999-82, RRID: AB_466125) as appropriate for the peptide for 15 minutes on ice, then fixed with 4% paraformaldehyde (PFA) for 15 minutes on ice. Cells were analyzed on a BD FACSymphony™ A5 SE Cell Analyzer.

Mean fluorescence intensity (MFI) values were normalized to the baseline and highest value within an experiment. Values were then plotted in GraphPad Prism (RRID: SCR_002798) and an S4PL equation with a Hill Slope of 1 was used to fit the data and generate IC50 values.

### Differential Scanning Fluorimetry Assay for pMHC Thermal Melting

MHC I monomers (NIH Tetramer Core) loaded with the UV-cleavable peptide FAPGNYJAL were combined with the exchange peptide in PBS at a concentration of 0.2 mg/mL monomer and 0.4 µM peptide in a 96 well plate. The solution was incubated for 1 hour in a UVA light box. Loaded monomers were centrifuged at 3300g for 5 minutes and supernatant transferred to a final tube. 18 µL of this solution was combined with 2 µL of 10X SYPRO Orange (Invitrogen, Cat. No. S6650) in a 96 well plate and incubated for 10 minutes to allow for the dye to bind the protein. The plate was assayed on a QuantStudio 7 Flex Real-Time PCR System; samples were equilibrated at 25°C for 1 minute then heated at a rate of 0.05°C/s from 25°C to 95°C with continuous fluorescence measurements at an excitation of 470 nm and emission of 570 nm. Melting curves were normalized to the minimum and maximum fluorescence value and the inverse of the first derivative (-dF/dT) was then calculated. The melting temperature is defined as the minimum value of -dF/dT.

### Bone Marrow-Derived Dendritic Cell Vaccination

Bone marrow cells were collected by euthanizing mice and removing the femur and tibia. The heads of the bones were clipped and placed facing down into a 0.5-mL tube nested in a 1.5-mL microcentrifuge tube. These tubes were flash spun at maximum speed in a microcentrifuge to extract the bone marrow. Red blood cells were lysed with 1X Red Blood Cell (RBC) Lysis Buffer (eBioscience; Cat. No. 00-4333-57). Cells were filtered through 70 µm cell strainers, counted and resuspended in Dendritic Cell Media (DCC) (DMEM, 10% FBS, 2 mM L-glutamine, 50 U/µL Pen/strep, 1X β-mercaptoethanol, 100 ng/mL Flt-3L and 3.75 ng/mL GM-CSF) at 1x10^6^ cells/mL. 1x10^7^ cells were plated per well in a 6 well plate and cultured at 37°C. On day 4, each well was supplemented with additional Flt-3L 1 µg, GM-CSF 37.5 ng and 0.5X β-mercaptoethanol. On day 7, the media and cells were collected, spun down at 1200 rpm for 5 minutes, resuspended in fresh DCC and replated 1x10^6^ cells/mL in a 6 well plate. On day 9, CpG (1 µg/mL) was added to generate mature bone marrow-derived dendritic cells (BMDCs). On day 10, the BMDCs were collected and resuspended in 10 mL of DCC media with 1 µg/mL of the appropriate neoantigen short-peptide in 15-mL conical tubes. These tubes were incubated with a loose cap in a 37°C incubator at 5% CO_2_ for 2 hours. The neoantigen-pulsed BMDCs were then washed with PBS and resuspended at 1.5x10^7^ cells/mL and 1.5 million cells were injected in 100 µL volume subcutaneously at the tail base. One week later, mice were boosted with 6 µg of neoantigen short-peptide and 25 µg of cyclic di-GMP adjuvant in PBS and injected in 100 µL volume subcutaneously at the tail base. Spleens were harvested 1 week after the booster dose.

Spleens were mashed through a 70 µm filter, spun down at 1200 rpm for 5 minutes, then incubated in 1X RBC Lysis Buffer for 5 minutes on ice. Pellets were then resuspended in R10 media (RPMI 1640, 10% FBS, 2 mM L-glutamine, 10 mM HEPES, 1X NEAA, 1X β-mercaptoethanol and 50 U/µL Penicillin/streptomycin). 1.5 million splenocytes were plated in R10 for 5 hours with 1X Brefeldin-A and 1 µg/mL of the corresponding neoantigen short-peptide. Cells were then washed and stained with Live/Dead Fixable Dead Cell Stain (Invitrogen, Cat. No. L34957) for 30 minutes on ice, then anti-mouse CD45 (Invitrogen, Cat. No. 47-0451-82, RRID: AB_1548781) and CD8*a* (BD Biosciences, Cat. No. 553031, RRID: AB_394569) antibodies for 15 minutes on ice. Cells were fixed for 1 hour at room temperature and permeabilized with the eBioscience Intracellular Fixation and Permeabilization Buffer Set (Invitrogen, Cat. No. 88-8824-00). Cells were then stained overnight with anti-mouse IFNγ (BioLegend, Cat. No. 505822, RRID: AB_961359) and TNF*a* (BioLegend, Cat. No. 506306, RRID: AB_315427) antibodies. Cells were analyzed on a BD FACSymphony™ A5 SE Cell Analyzer.

### Analysis of Public Training Data

Training data from MHCflurry2.0 was imported and filtered to only contain quantitative affinity measurements. These 170,722 entries were sorted for duplicates with matched MHC and peptide identities; there were a total of 11,645 MHC-peptide combinations that had more than one entry, creating a total of 14,577 pairs. Many of these pairs were duplicate entries of the same measurement, typically arising when compiled affinity tables were re-entered into IEDB. We removed these duplicated pairs by filtering out measurements that fell within 3% of each other. Additionally, peptides whose measurements were at the limit of detection, were removed from the visualizations, for a total of 4,174 pairs.

Analyses were performed in Python 3.12.2 (RRID: SCR_008394) with numpy 2.5.1 (RRID: SCR_008633), pandas 3.0.5 (RRID: SCR_018214), scipy 1.18.0 (RRID: SCR_008058) and matplotlib 3.11.1 (RRID: SCR_008624).

The datasets generated in this study are available from the corresponding author upon reasonable request. These include flow cytometry data from the RMA-S affinity assays and in vivo vaccination experiments, as well as the raw melting curve values.

## ACKNOWLEDGEMENTS

We would like to acknowledge S.L. Shanahan for supplying the RMA-S cell line; N. Struntz for input on modeling IC50 curves; P. Canaday from the Flow Cytometry and Monoclonal Antibody Core for her technical assistance. We thank the NIH Tetramer Core Facility (contract number 75N93020D00005) for providing the UV-exchangeable monomers.

This work was supported by a V Scholar Grant (V2023-012) from the V Foundation for Cancer Research, an R37 from the NIH National Cancer Institute (R37CA302926), a Lung Cancer Discovery Award (LCD-1034504) from the American Lung Association and a pilot grant from the OHSU Brenden-Colson Center for Pancreatic Care to M.L.B., and a Frohnmayer Hicks Sciarretta Cancer Research Scholars Program fellowship to P.J.M. This work utilized the OHSU Flow Cytometry and Monoclonal Antibody Shared Resource (RRID:SCR_009974) and the OHSU Biostatistics Shared Resource, supported by the Knight Cancer Institute NCI Comprehensive Cancer Center grant (P30CA069533).

## AUTHOR CONTRIBUTIONS

P.J.M., M.L.B. and M.K.R designed the study; P.J.M., C.N.S., V.P.S., A.M.G., R.B.C. and B.M.M. performed the experiments; P.J.M. and M.L.B. wrote the manuscript with input from other authors.

## REFERENCES

1. Sharma, P. et al. Immune checkpoint therapy—current perspectives and future directions. Cell 186, 1652–1669 (2023).

2. Puig-Saus, C. et al. Neoantigen-targeted CD8+ T cell responses with PD-1 blockade therapy. Nature 615, 697–704 (2023).

3. Simoni, Y. et al. Bystander CD8+ T cells are abundant and phenotypically distinct in human tumour infiltrates. Nature 557, 575–579 (2018).

4. Oliveira, G. et al. Phenotype, specificity and avidity of antitumour CD8+ T cells in melanoma. Nature 596, 119–125 (2021).

5. Lybaert, L. et al. Neoantigen-directed therapeutics in the clinic: where are we? Trends in Cancer 9, 503–519 (2023).

6. Merck and Moderna Announce Phase 3 INTerpath-001 Trial of Intismeran Autogene Plus KEYTRUDA® Met Endpoints of Recurrence-Free Survival (RFS) and Distant Metastasis-Free Survival (DMFS) in Patients With Completely Resected Stage IIB-IV Melanoma. Merck.com https://www.merck.com/news/merck-and-moderna-announce-phase-3-interpath-001-trial-of-intismeran-autogene-plus-keytruda-met-endpoints-of-recurrence-free-survival-rfs-and-distant-metastasis-free-survival-dmfs-in-patient/.

7. Barnett, K. K. et al. FDA Approval Summary: Afamitresgene Autoleucel for Adults with HLA-Restricted, MAGE-A4–Positive Unresectable or Metastatic Synovial Sarcoma after Prior Chemotherapy. Clin Cancer Res 31, 3112–3117 (2025).

8. DuPage, M. et al. Endogenous T cell responses to antigens expressed in lung adenocarcinomas delay malignant tumor progression. Cancer Cell 19, 72–85 (2011).

9. Mellman, I., Chen, D. S., Powles, T. & Turley, S. J. The cancer-immunity cycle: Indication, genotype, and immunotype. Immunity 56, 2188–2205 (2023).

10. Roerden, M. et al. Neoantigen architectures define immunogenicity and drive immune evasion of tumors with heterogenous neoantigen expression. J Immunother Cancer 12, e010249 (2024).

11. Burger, M. L. et al. Antigen dominance hierarchies shape TCF1+ progenitor CD8 T cell phenotypes in tumors. Cell 184, 4996–5014.e26 (2021).

12. Antsiferova, M. et al. Novel immunodominant neoepitope in a KPC mouse model of pancreatic cancer allowing identification of tumor-specific T cells. Oncoimmunology 14, 2489815.

13. Kim, S. I. et al. CD4+ T Cells Mediate MHC-Deficient Tumor Rejection and Endothelial Cell Reprogramming. Cancer Immunol Res 14, 107–121 (2026).

14. Yewdell, J. W. & Bennink, J. R. Immunodominance in major histocompatibility complex class I-restricted T lymphocyte responses. Annu Rev Immunol 17, 51–88 (1999).

15. Wellington, D., Yin, Z., Kessler, B. M. & Dong, T. Immunodominance complexity: lessons yet to be learned from dominant T cell responses to SARS-COV-2. Curr Opin Virol 50, 183–191 (2021).

16. Anikeeva, N., Gakamsky, D., Schøller, J. & Sykulev, Y. Evidence that the Density of Self Peptide-MHC Ligands Regulates T-Cell Receptor Signaling. PLoS One 7, e41466 (2012).

17. Harndahl, M. et al. Peptide-MHC class I stability is a better predictor than peptide affinity of CTL immunogenicity. Eur J Immunol 42, 1405–1416 (2012).

18. Jurtz, V. et al. NetMHCpan 4.0: Improved peptide-MHC class I interaction predictions integrating eluted ligand and peptide binding affinity data. J Immunol 199, 3360–3368 (2017).

19. O’Donnell, T. J., Rubinsteyn, A. & Laserson, U. MHCflurry 2.0: Improved Pan-Allele Prediction of MHC Class I-Presented Peptides by Incorporating Antigen Processing. Cell Syst 11, 42–48.e7 (2020).

20. Bonsack, M. et al. Performance Evaluation of MHC Class-I Binding Prediction Tools Based on an Experimentally Validated MHC–Peptide Binding Data Set. Cancer Immunol Res 7, 719–736 (2019).

21. Paul, S. et al. Benchmarking predictions of MHC class I restricted T cell epitopes in a comprehensively studied model system. PLoS Comput Biol 16, e1007757 (2020).

22. AI-driven computational methods and benchmarking for T-cell antigen identification | Briefings in Bioinformatics | Oxford Academic. https://academic.oup.com/bib/article/27/2/bbag123/8526139.

23. Wolf, Y. et al. UVB-Induced Tumor Heterogeneity Diminishes Immune Response in Melanoma. Cell 179, 219–235.e21 (2019).

24. Westcott, P. M. K. et al. Mismatch repair deficiency is not sufficient to elicit tumor immunogenicity. Nat Genet 55, 1686–1695 (2023).

25. Capietto, A.-H. et al. Mutation position is an important determinant for predicting cancer neoantigens. J Exp Med 217, e20190179 (2020).

26. Li, S. et al. Characterization of neoantigen-specific T cells in cancer resistant to immune checkpoint therapies. Proc Natl Acad Sci U S A 118, e2025570118 (2021).

27. Yadav, M. et al. Predicting immunogenic tumour mutations by combining mass spectrometry and exome sequencing. Nature 515, 572–576 (2014).

28. Hos, B. J. et al. Identification of a neo-epitope dominating endogenous CD8 T cell responses to MC-38 colorectal cancer. Oncoimmunology 9, 1673125 (2019).

29. Nimanong, S. et al. CD40 Signaling Drives Potent Cellular Immune Responses in Heterologous Cancer Vaccinations. Cancer Res 77, 1918–1926 (2017).

30. Gubin, M. M. et al. Checkpoint Blockade Cancer Immunotherapy Targets Tumour-Specific Mutant Antigens. Nature 515, 577–581 (2014).

31. Gebert, J. et al. Recurrent Frameshift Neoantigen Vaccine Elicits Protective Immunity With Reduced Tumor Burden and Improved Overall Survival in a Lynch Syndrome Mouse Model. Gastroenterology 161, 1288–1302.e13 (2021).

32. Howarth, M., Williams, A., Tolstrup, A. B. & Elliott, T. Tapasin enhances MHC class I peptide presentation according to peptide half-life. Proceedings of the National Academy of Sciences 101, 11737–11742 (2004).

33. van der Most, R. G. et al. Changing immunodominance patterns in antiviral CD8 T-cell responses after loss of epitope presentation or chronic antigenic stimulation. Virology 315, 93–102 (2003).

34. Udaka, K., Wiesmüller, K. H., Kienle, S., Jung, G. & Walden, P. Self-MHC-restricted peptides recognized by an alloreactive T lymphocyte clone. J Immunol 157, 670–678 (1996).

35. Niedermann, G. et al. Contribution of proteasome-mediated proteolysis to the hierarchy of epitopes presented by major histocompatibility complex class I molecules. Immunity 2, 289–299 (1995).

36. Van Der Burg, S. H., Visseren, M. J., Brandt, R. M., Kast, W. M. & Melief, C. J. Immunogenicity of peptides bound to MHC class I molecules depends on the MHC-peptide complex stability. The Journal of Immunology 156, 3308–3314 (1996).

37. Busch, D. H. & Pamer, E. G. MHC class I/peptide stability: implications for immunodominance, in vitro proliferation, and diversity of responding CTL. J Immunol 160, 4441–4448 (1998).

38. Müllbacher, A. et al. High peptide affinity for MHC class I does not correlate with immunodominance. Scand J Immunol 50, 420–426 (1999).

39. Frontiers | NetMHCpan-4.2: improved prediction of CD8+ epitopes by use of transfer learning and structural features. https://www.frontiersin.org/journals/immunology/articles/10.3389/fimmu.2025.1616113/full.

40. Reynisson, B., Alvarez, B., Paul, S., Peters, B. & Nielsen, M. NetMHCpan-4.1 and NetMHCIIpan-4.0: improved predictions of MHC antigen presentation by concurrent motif deconvolution and integration of MS MHC eluted ligand data. Nucleic Acids Research 48, W449–W454 (2020).

41. Andreatta, M. & Nielsen, M. Gapped sequence alignment using artificial neural networks: application to the MHC class I system. Bioinformatics 32, 511–517 (2016).

42. Kim, Y., Sidney, J., Pinilla, C., Sette, A. & Peters, B. Derivation of an amino acid similarity matrix for peptide: MHC binding and its application as a Bayesian prior. BMC Bioinformatics 10, 394 (2009).

43. Peters, B. & Sette, A. Generating quantitative models describing the sequence specificity of biological processes with the stabilized matrix method. BMC Bioinformatics 6, 132 (2005).

44. Zhang, H., Lund, O. & Nielsen, M. The PickPocket method for predicting binding specificities for receptors based on receptor pocket similarities: application to MHC-peptide binding. Bioinformatics 25, 1293–1299 (2009).

45. Karosiene, E., Lundegaard, C., Lund, O. & Nielsen, M. NetMHCcons: a consensus method for the major histocompatibility complex class I predictions. Immunogenetics 64, 177–186 (2012).

46. Schumacher, T. N. et al. Direct binding of peptide to empty MHC class I molecules on intact cells and in vitro. Cell 62, 563–567 (1990).

47. Hellman, L. M. et al. Differential scanning fluorimetry based assessments of the thermal and kinetic stability of peptide-MHC complexes. J Immunol Methods 432, 95–101 (2016).

48. Peters, B. et al. A Community Resource Benchmarking Predictions of Peptide Binding to MHC-I Molecules. PLoS Comput Biol 2, e65 (2006).

49. Vita, R. et al. The Immune Epitope Database (IEDB): 2024 update. Nucleic Acids Res 53, D436–D443 (2024).

50. Bloom, M. B. et al. Identification of Tyrosinase-related Protein 2 as a Tumor Rejection Antigen for the B16 Melanoma. J Exp Med 185, 453–460 (1997).

51. McCarron, M. J. et al. Cross-competition shapes CD8+ T cell hierarchies and differentiation after RNA vaccination. 2025.10.26.684631 Preprint at 10.1101/2025.10.26.684631 (2025).

52. Alspach, E. et al. MHC-II neoantigens shape tumour immunity and response to immunotherapy. Nature 574, 696–701 (2019).

53. Huang, S. et al. A Programmable Nanovaccine Platform Based on M13 Bacteriophage for Personalized Cancer Vaccine and Therapy. Adv Mater 37, e10229 (2025).

54. Strønen, E. et al. Targeting of cancer neoantigens with donor-derived T cell receptor repertoires. Science 352, 1337–1341 (2016).

55. Roerden, M. & Spranger, S. Cancer immune evasion, immunoediting and intratumour heterogeneity. Nat Rev Immunol 25, 353–369 (2025).

56. Duan, F. et al. Genomic and bioinformatic profiling of mutational neoepitopes reveals new rules to predict anticancer immunogenicity. J Exp Med 211, 2231–2248 (2014).

57. Freed-Pastor, W. A. et al. The CD155/TIGIT axis promotes and maintains immune evasion in neoantigen-expressing pancreatic cancer. Cancer Cell 39, 1342–1360.e14 (2021).

58. Schrörs, B. et al. MC38 colorectal tumor cell lines from two different sources display substantial differences in transcriptome, mutanome and neoantigen expression. Front Immunol 14, 1102282 (2023).

59. Kessler, J. H. et al. Antigen processing by nardilysin and thimet oligopeptidase generates cytotoxic T cell epitopes. Nat Immunol 12, 45–53 (2011).

60. Hammer, G. E., Gonzalez, F., James, E., Nolla, H. & Shastri, N. In the absence of aminopeptidase ERAAP, MHC class I molecules present many unstable and highly immunogenic peptides. Nat Immunol 8, 101–108 (2007).

61. Van den Eynde, B. J. & Morel, S. Differential processing of class-I-restricted epitopes by the standard proteasome and the immunoproteasome. Curr Opin Immunol 13, 147–153 (2001).

62. Jenkins, M. K. & Moon, J. J. The role of naïve T cell precursor frequency and recruitment in dictating immune response magnitude. J Immunol 188, 4135–4140 (2012).

63. Rojas, L. A. et al. Personalized RNA neoantigen vaccines stimulate T cells in pancreatic cancer. Nature 618, 144–150 (2023).

64. Li, S. et al. Bystander CD4+ T cells infiltrate human tumors and are phenotypically distinct. Oncoimmunology 11, 2012961 (2022).

65. Holm, J. S. et al. Neoantigen-specific CD8 T cell responses in the peripheral blood following PD-L1 blockade might predict therapy outcome in metastatic urothelial carcinoma. Nat Commun 13, 1935 (2022).

66. Braun, D. A. et al. A neoantigen vaccine generates antitumour immunity in renal cell carcinoma. Nature 639, 474–482 (2025).

67. Ott, P. A. et al. An immunogenic personal neoantigen vaccine for patients with melanoma. Nature 547, 217–221 (2017).

68. Hu, Z. et al. Personal neoantigen vaccines induce persistent memory T cell responses and epitope spreading in patients with melanoma. Nat Med 27, 515–525 (2021).

69. Rappaport, A. R. et al. A shared neoantigen vaccine combined with immune checkpoint blockade for advanced metastatic solid tumors: phase 1 trial interim results. Nat Med 30, 1013–1022 (2024).

